# Bayesian Prior Uncertainty and Surprisal Elicit Distinct Neural Patterns During Sound Localization in Dynamic Environments

**DOI:** 10.1101/2024.07.22.604566

**Authors:** Burcu Bayram, David Meijer, Roberto Barumerli, Michelle Spierings, Robert Baumgartner, Ulrich Pomper

**Affiliations:** Department of Cognition, Emotion, and Methods in Psychology, Faculty of Psychology, University of Vienna, Vienna, Austria; Austrian Academy of Sciences, Acoustics Research Institute, Vienna, Austria; Department of Behavioral and Cognitive Biology, University of Vienna, Vienna, Austria; Department of Behavioral Biology, Leiden University, Leiden, Netherlands

**Author notes:** corresponding author: Burcu Bayram.

**Keywords:** Perception, Bayesian inference, Neural oscillations, Auditory, Localization, EEG, Prior uncertainty, Surprisal

## Abstract

Estimating the location of a stimulus is a key function in sensory processing, and widely considered to result from the integration of prior information and sensory input according to Bayesian principles. A deviation of sensory input from the prior elicits surprisal, depending on the uncertainty of the prior.

While this mechanism is increasingly understood in the visual domain, much less is known about its implementation in audition, especially regarding spatial localization. Here, we combined human EEG with computational modeling to study auditory spatial inference in a noisy, volatile environment and analyzed behavioral and neural patterns associated with prior uncertainty and surprisal.

First, our results demonstrate that participants indeed used prior information during periods of stable environmental statistics, but showed evidence of surprisal and discarded prior information following environmental changes. Second, we observed distinct EEG activity patterns associated with prior uncertainty and surprisal in both the time- and time-frequency domain, which are in line with previous studies using visual tasks. Third, these EEG activity patterns were predictive of our participants’ sound localization error, response uncertainty, and prior bias on a trial-by-trial basis.

In summary, our work provides novel behavioral and neural evidence for Bayesian inference during dynamic auditory localization.

## Introduction

In stable environments, perception can benefit from past experiences, especially when our sensory representations are unreliable. Here, a mismatch between prior and sensory input results in prediction error, which can be used to update predictions and increase perceptual accuracy. However, following an abrupt change in the environment, prior information quickly becomes irrelevant or even detrimental for the perceptual decision-making process^1,2^. One way to conceptualize optimal decision making in such dynamic environments is via Bayesian inference, in which perception is based on the integration of prior knowledge and new sensory observations, weighted by their reliabilities and the inferred probability of an environmental change^3,4^.

In the past, many studies have examined Bayesian inference using change-point paradigms, in which a non-stationary environment is simulated by pseudo-randomly changing stimulus statistics throughout the experimental task^4,5^. In such dynamic environments, changes in the environment lead to large surprisal and indicate that the prior is not relevant anymore. Thereby, humans have been shown to perform similar to an ideal Bayesian observer in settings such as visual spatial localization^6,7^, visual orientation discrimination^8^, and auditory pitch discrimination^9^, although sub-optimality in decision making has also been reported^10–12^. For instance, participants might not weight the sensory evidence and prior information according to their reliabilities^12^, especially when the task complexity is high^10^. At the neural level, a number of studies have found prediction signals (i.e. the prior) reflected in beta-band (14 – 30 Hz)^13–15^ and surprisal reflected in gamma-band (40-100 Hz) oscillatory activity^14–16^. Sedley and colleagues^15^ employed a pitch discrimination task to disentangle neural patterns associated with surprisal and prediction precision (i.e. inverse of prior uncertainty). They observed that surprisal was reflected in gamma-band oscillations starting from ∼250 ms post-stimulus, and prior uncertainty about the next stimulus was positively correlated with alpha-band oscillations (8-12 Hz) starting from ∼280 ms post-stimulus. This suggests that the prior, its precision and surprisal are coded in distinct neural patterns. Similarly, Chao et al.^14^ used a hierarchical predictive coding model to differentiate the feedforward and feedback signals during a tone sequence discrimination task. They observed feedback prediction signals in beta range oscillations during the pre-stimulus time period, and feedforward prediction error signals reflected in gamma-band oscillations following stimulus presentation. From a predictive coding perspective^17^, the above studies demonstrate that surprisal (or prediction errors) and prior are coded in higher (i.e. gamma) and lower (beta) frequency bands, respectively, while uncertainty associated with the prior information is reflected in alpha-band oscillations. In addition to gamma-band oscillations, surprisal is commonly observed to scale with the amplitude of the P3 event-related potential (ERP), along with the updating of the prior information (i.e. belief update)^8,18–20^. Nassar and colleagues^8^ found that the effect of P3 amplitude and surprisal on belief update was dependent on the statistical context in the environment. They used a change-point paradigm in which larger surprisal values would indicate a true change in the environment as well as an oddball paradigm in which the oddball stimuli would trigger large surprisal responses without a change in the environment. Their results showed that contrary to the change-point paradigm, participants did not update their beliefs even with high surprisal values in response to oddball stimuli, even though in both conditions P3 amplitude scaled with surprisal. So far, evidence for a Bayesian inference mechanism comes primarily from the visual domain, as well as from auditory pitch or temporal estimation tasks^6,15,20^. However, Krishnamurthy et al.^21^ have provided an intriguing behavioral study on Bayesian integration of prior information and sensory evidence in auditory spatial localization. In their task, participants had to first predict and then estimate the actual location of a sound source, as the predictability of its location varied over time. They observed that changes in stimulus predictability lead to changes in the magnitude of prior-driven biases, dependent on the relevance and reliability of prior expectations. In other words, periods of stable stimulus statistics enhanced prior usage, while recent changes reduced it. However, their study provided an additional visual representation for all the sounds establishing the prior, which arguably leads to multisensory (i.e. audio-visual) rather than purely auditory spatial priors. As humans perform better at localization of visual compared to auditory stimuli in unimodal tasks^22,23^, vision is usually the dominant modality during multisensory localization^24,25^. In addition, multisensory recalibration effects can transfer to subsequent unimodal tasks^26^ and subjects trained with audio-visual stimuli are more accurate in their sound localization responses compared to subjects trained using only auditory stimuli^27^, while, modality dominance is reduced or even reversed with decreasing reliability of the visual stimuli^28,29^. For these reasons, it is unclear to what extend the results of Krishnamurthy et al.^21^ specifically reflect Bayesian inference in auditory spatial localization, or are at least partly driven by visual priors. Consequently, the aim of our present study is to expand upon the literature by (1) investigating whether behavioral responses in a unimodal auditory localization task adhere to the principles of Bayesian inference; (2) testing how this spatial inference process is altered by the presence of additional visual priors (as in Krishnamurthy et al.^21^); (3) study the neural patterns associated with prior uncertainty and surprisal, along with their impact on subsequent behavioral responses.

Participants listened to sequences of sounds coming from pseudorandom locations and reported the location of the last sound of each trial. We conducted both an audio-visual condition (as in Krishnamurthy et al.^21^) and a modified audio-only condition while recording high density EEG, and fitted a near optimal Bayesian observer model to participants’ responses. Our results reveal neural patterns reflecting prior uncertainty and surprisal for both conditions in time-domain as well as the lower (i.e. alpha/beta) and higher (i.e. gamma) frequency range oscillations. Critically, we observed a significant relationship between the neural activity associated with prior uncertainty and surprisal and the behavioural location estimation error, response uncertainty, and prior bias. In summary, our results indicate behaviourally relevant electrophysiological patterns reflecting Bayesian inference processes during both auditory-only and audio-visual spatial localization in dynamic environments.

## Materials and Methods

### Participants

Thirty-five participants took part in the study, in exchange for monetary compensation (10 Euros per hour). Our sample size is based on a previous study employing a similar auditory change-point paradigm^21^. Three participants had a mean estimation error of > 25° during the training sessions and were excluded prior to the main experiment. We excluded three additional participants due to a large number of noisy EEG channels (> 10%). The remaining 29 participants were between 19 and 37 years old (15 females, M_age_ = 24.4, SD_age_ = 3.8) and right-handed. They reported no hearing impairments or neurological deficits, had normal or corrected-to-normal vision, gave informed consent, and were naïve to the purpose of the experiment. The study was conducted in accordance with the standards of the Declaration of Helsinki (1996). We further followed the Austrian Universities Act of 2002, which states that only medical universities or studies conducting applied medical research are required to obtain additional approval by an ethics committee. Therefore, no additional ethical approval was required for our study.

### Experimental setup and procedure

The experiment was run using MATLAB (2018b, MathWorks, Natick, MA) and Psychophysics Toolbox^30^. We presented visual stimuli via an LCD monitor (48 x 27 cm) with a refresh rate of 60 Hz and auditory stimuli via tube earphones (ER2; Etymotic Research, Elk Grove Village, IL). Individual auditory stimuli consisted of 50 ms pink noise burst (10 ms on- and off-set ramps), high-pass filtered using a 4^th^ order Butterworth filter with a 250 Hz cut-off frequency. Each sound was rendered at a specific direction by employing each participant’s head-related transfer function (HRTF, measured prior to the experimental sessions, using the same approach as in Ignatiadis et al.^31^) using the Auditory Modeling Toolbox^32^.

Low visual and auditory stimulus presentation latencies were verified via an oscilloscope (M = 0 ms, SD = 4.4 ms). Participants sat at a desk in a dark and sound attenuated room, with their head in a chin rest (75 cm distance to the screen) to minimize movement.

The experiment was divided into two sessions, conducted on separate days (less than 3 days apart). The first session consisted of nine training blocks (not included in our analysis), to familiarize participants with the sound localization task and response method. In the second session, participants completed two further training blocks and six main task blocks. Each block consisted of 50 trials, resulting in 300 trials for the main task.

Throughout the main task, we collected 128-channel high-density EEG (actiCAP with actiCHamp; Brain Products GmbH, Gilching, Germany) and eye tracking data (EyeLink 1000 Plus; SR Research, Osgoode, Ontario, Canada), at a sampling rate of 1 kHz. Electrode impedances were kept below 25 kΩ and the signal was recorded against ‘FCz’ as reference electrode. In addition to the scalp electrodes, we recorded an auxiliary audio channel via a stimulus tracking device (StimTrak, Brain Products GmbH, Gilching, Germany) to later align the EEG triggers offline with the onset of sounds and ensure correct trigger timing.

### Training task

At the beginning of each trial, participants fixated on a central dot (0.5° radius) displayed at the centre of the screen. Around the fixation dot, we displayed a semi-arc (0.75° width) that represented the range of azimuth angles on the frontal horizontal plane (see Figure 1). After 750-1000 ms of fixation, jittered on each trial, we presented a single sound from different locations in azimuth (from 90° to -90°). 950 ms after the sound offset, a mouse cursor appeared at the location of the fixation dot, which signalled participants to respond and turned into a line when moved close to the semi-arc. Their task then was to indicate the azimuth location of the sound, by moving the line along the semi-arc (response resolution <1°) via a mouse in their right hand, and click the left mouse button at the desired location. Subsequently, they indicated their level of response uncertainty by marking an 80% confidence-interval around their response location. After that, feedback was provided for 500 ms via a red line presented on the arc at the true sound location, followed by a new trial. At the end of each training block, participants received additional feedback on their mean absolute error (in degrees) and the percentage of times that their marked area of response uncertainty included the true sound location (goal was 80%).

**Figure 1.**
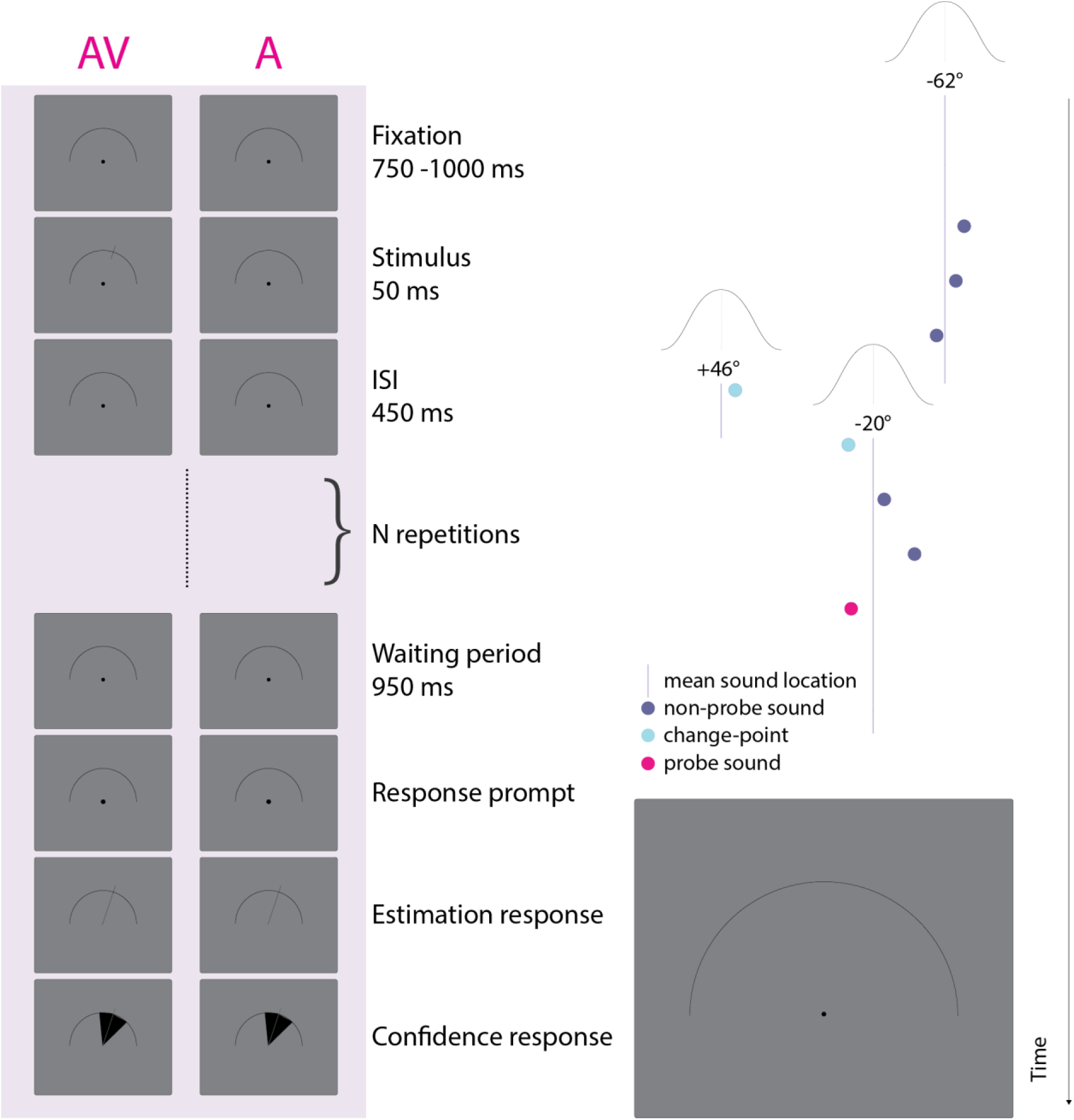
Experimental design. Left part: An exemplary trial depicted for both audio-visual (AV) and audio-only (A) conditions. Each trial starts with a fixation period followed by a sequence of sounds. In the A condition, only auditory stimuli are presented. In the AV condition, the respective sound location is simultaneously shown via a line on the semi-arc (for all sounds except for the final probe sound). At the end of the sequence, participants indicate the location of the probe sound by rotating the response line on the semi-arc. Subsequently, they mark an 80% confidence interval around the response line. Right part: Depiction of the sampling of sound locations for an exemplary trial. Sound locations are sampled from a Gaussian distribution with standard deviation of 10° (i.e. experimental noise). Along with each change-point sound (turquoise circle), a new mean sound location is sampled. Four sounds are presented following the last change-point in the sequence and therefore, the trial has a stimulus-after-change-point (SAC) level of 4.

Importantly, trials alternated between an *auditory-only* (A) (as described above) and an *audio-visual* (AV) condition. In the AV condition, a line indicating the sound location appeared on the arc simultaneously with the sound presentation. We added this condition to the training, to help participants establish a mapping between the auditory sound location and its abstract spatial representation on the visual arc on screen.

### Main task

The main task was highly similar to the training, with the important difference that each trial consisted of a sequence of sounds, with a stimulus-onset asynchrony of 500 ms. Sound locations were randomly sampled from a normal distribution whose (generative) mean was sampled from a uniform distribution bounded between 60 and -60 degrees, and a constant standard deviation of 10° (i.e. experimental noise). Each sound of the sequence had a 1/6 probability of being a change-point, at which point a new generative mean was sampled from the bounded uniform distribution (60° to -60°), thus resulting in a sudden change of the mean sound location.

The participants’ task was to indicate the location of the last sound of the sequence (i.e. the probe sound). Each sequence contained between 1 and 43 sounds and each sound had a 1/12 probability of being the probe sound, thus rendering the trial length unpredictable and encouraging participants to pay attention to each sound location. Prior to the experiment, participants were only superficially informed about the concepts of change-points and experimental noise, without mentioning specific parameter values.

Akin to the training blocks, we presented the A and AV condition trials in alternating order. In AV trials, all sounds except for the final probe sound were presented along with a simultaneous visual representation of the sound location via a line on the semi-arc. To ensure comparability of the two conditions, we presented identical trials in the A and AV conditions. In other words, each A trial had a corresponding AV trial, which only differed in the additional visual representations for the latter. The order of trials was randomized within subjects, and the corresponding A and AV trials were never presented in direct succession.

As in the training task, participants gave localization and response uncertainty responses at the end of each trial. However, they did not receive immediate feedback on the true sound location after each trial, but only a summary performance feedback at the end of each block, regarding their mean absolute error (in degrees) and the percentage of times that their marked area of response uncertainty included the true sound location.

### Bayesian model

We fitted a near optimal Bayesian observer model to our participants’ localization responses (for modeling details see Supplement 1) to obtain two latent variables (i.e. not directly observable variables, which we inferred via our model) for every stimulus with optimized parameters for each participant. The first latent variable is the prior uncertainty (*PU*), which we defined as standard deviation of the preceding posterior distribution (see Supplement 1, Eq. 25). The second latent variable is the information theoretic quantity of surprisal (*SU*). This is defined as the negative logarithm of the probability density of the full prior distribution, evaluated at the latest observation (see Supplement 1, Eq. 26)

### Behavioural analysis

Using the participants’ responses, we derived three behavioural metrics for each trial and analysed them over two experimental factors. The behavioural metrics are: (i) the *estimation error,* which is calculated by computing the absolute difference between the localization response and the sound’s true location;.(ii) the *response uncertainty,* indicating the confidence area (in degrees) that participants marked around their response line at the end of each trial; (iii) the *prior bias,* computed as the estimation error on the current trial, divided by the difference (in degrees) between the previous and the current sound locations:

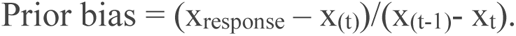

These metrics were evaluated over two main independent variables. The first one is the sensory modality condition (AV vs. A), the second one is how many individual sounds had been presented since the last change-point (sounds after change-point, SAC). In other words, the latter is a proxy of the strength of the present prior and the expected surprise elicited by the current sound.

To investigate how our experimental manipulations affected the behavioral variables, we computed a linear mixed effect model (LMM) using the glmmTMB^33^ package in R (version 4.3.2) with Modality (AV and A) and SAC level (1 to 6) as fixed effect variables and subjects as random effect variable. Single sound trials are excluded from the analyses.

### EEG preprocessing

All EEG data preprocessing and analysis was performed using EEGlab^34^ (version 2022.1), Fieldtrip^35^ (version 20220827), and custom scripts.

First, we aligned our EEG triggers to sound onsets using the Stimtrak external channel. Next, we downsampled the continuous EEG signal to 250 Hz, applied a high-pass filter with a cut-off frequency of 0.25 Hz (Hamming window, zero-phase finite impulse response filter) using the ‘*pop_eegfiltnew*’ function from EEGlab and removed line noise using the ‘*nt_zapline*’ function from NoiseTools toolbox^36^. We then epoched the data from -1 to 2 seconds relative to the onset of each sound stimulus in the experiment, visually inspected the data and excluded epochs containing excessive noise as well as eye blinks close to the stimulus onset. As mentioned above, identical trials were presented in the A and AV conditions, although in randomized order. To maximize comparability between the A and AV conditions, for each participant, we only analyzed trial epochs that were present in both conditions after artifact rejection. Therefore, if we rejected an epoch in one condition, we also rejected the corresponding epoch in the other condition.

We then interpolated noisy channels via spherical interpolation, added the reference electrode ‘FCz’ back to the dataset and re-referenced the data to the average of all electrodes.

Next, we performed independent component analysis (ICA) to identify and remove ocular and heart-related artifacts. Particularly, in order to avoid using overlapping data segments for the ICA we performed the ICA decomposition on slightly differently preprocessed data: We high-pass filtered the data at 1 Hz cut-off frequency^37^ and extracted the epochs between 0 to 500 ms for each sound (thus containing no overlapping data periods between epochs). The remaining preprocessing steps were identical to the original dataset. We obtained ICA weights from this alternatively preprocessed dataset using the Picard algorithm with PCA (principal component analysis) to control for data rank deficiency caused by channel interpolation. Then we applied the resulting ICA weights back to the original dataset and rejected ocular and heart-related components by visual inspection using the IClabel^38^ plugin. Finally, we baseline corrected each epoch using the time period from -100 to 0 ms, relative to the sound onset.

### Spectral analysis

Prior to the spectral analysis, we subtracted the mean activity of all trials from each trial in to analyze only the induced oscillatory power. For the lower frequency range (4 to 30 Hz), we performed short-time fast Fourier transform, using a single Hanning taper with a window length of 300 ms in steps of 16 ms and a frequency resolution of 1 Hz. For the higher frequency range (40 to 100 Hz), we used the multi-taper method, with varying window length of 250-100 ms (window length shortens with higher frequencies) in steps of 16 ms and a frequency resolution of 2 Hz. The resulting spectral power was then expressed as relative signal change to the mean of the time period from 0 to 500 ms around each sound (i.e. the entire interval between the onset of the current and the next sound).

### Regression analysis

Next, our goal was to identify neural activity patterns associated with our Bayesian model latent variables PU (prior uncertainty) and SU (surprise). To do so, we performed three separate ordinary least squares linear regressions with both PU and SU values as predictors and EEG amplitudes as the output: one for the time-domain data (i.e. ERPs), one for the lower, and one for the higher frequency range. For the time-domain data, independent regressions were performed for each EEG time-point and channel, while for the time-frequency domain data, regressions also included the frequency domain.

Both EEG data and latent variables were z-transformed prior to the regression. We excluded the final probe sounds of each sequence from this analysis, since they contain only auditory stimuli in both A and AV conditions. Additionally, not including the final sounds that participants responded upon in the regression analysis, allowed us to keep them as an independent data set to test the behavioral relevance of the regression results (see next subsection). The first sounds were also excluded from the analysis, as they do not have a reliable estimate of the model latent variable values considering there are no preceding sounds to form a prior.

For each subject, this analysis resulted in a beta coefficient per time-point, channel, and frequency (for the time-frequency domain data), separately for each latent variable. For group statistics, we performed a total of six cluster-corrected permutation tests^39^, one for each combination of latent variable (PU, SU) and EEG data (time domain, low – and high time-frequency domain), to compare whether the obtained beta coefficients differed significantly from zero across participants. Cluster correction for multiple comparisons was applied using a cluster-level alpha of .001, and permutation of the data points was performed over 1000 iterations, and we only considered clusters with a duration of > 5ms length. For PU, we tested the time period of -250 ms to 100 ms relative to every sound onset, as the prior uncertainty should be neurally represented already pre-stimulus, and up until completion of initial sensory processing. For SU, on the other hand, we ran the test for the time period of 0 to 500 ms around each sound onset, as meaningful neural correlates of surprisal should only appear following stimulus presentation.

Additionally, we tested how the neural activity patterns associated with PU and SU variables differed between the AV and A conditions. Again, we used a cluster-based permutation test with the cluster level alpha set to .001 and permutation of the data points performed over 1000 iterations.

### Behavioral correlates of latent variable brain activation patterns

Finally, we were interested in whether the neural activity patterns associated with PU and SU, as identified in the previous step, were predictive of the behavioral metrics on a trial-by-trial basis. Therefore, within each subject, we regressed EEG activity around the final probe sound of each trial against the three behavioral variables estimation error, response uncertainty, and prior bias. Specifically, we averaged the EEG activity around the probe sound across all constituting data points (time, channels, and frequencies) for every significant PU and SU regression cluster. This resulted in a single value per trial, which we then regressed against the behavioral outcome of that trial (all variables were z-transformed prior to regression). For the group-level statistic, we performed one sample t-test (α = 0.05) for each regression between EEG activity and behavioral outcome variables, to see whether the resulting beta coefficients differed significantly from zero. Single sound trials are excluded from the analyses.

## Results

### Behavioral results

Figure 2 shows the behavioral and model data as a function of our two main independent variables, stimulus condition (AV vs. A) and SAC level (1 to 6). For the behavioral data, the linear mixed effects model revealed a smaller sound localization estimation error in the AV compared to the A condition (*p* < .01; see Table 1 for detailed statistical outcomes), as well as a larger prior bias (*p* < .001). In line with our expectations, these results suggest that the additional visual stimulus during non-probe sounds helped establishing a prior that eventually improved probe localization. Further, we observed a main effect of SAC level (*p* < .01) for prior bias, indicating that bias was small immediately following a change-point, but increased subsequently. No other main effects or interactions were observed for the behavioral data (*p* > .05).

**Figure 2.**
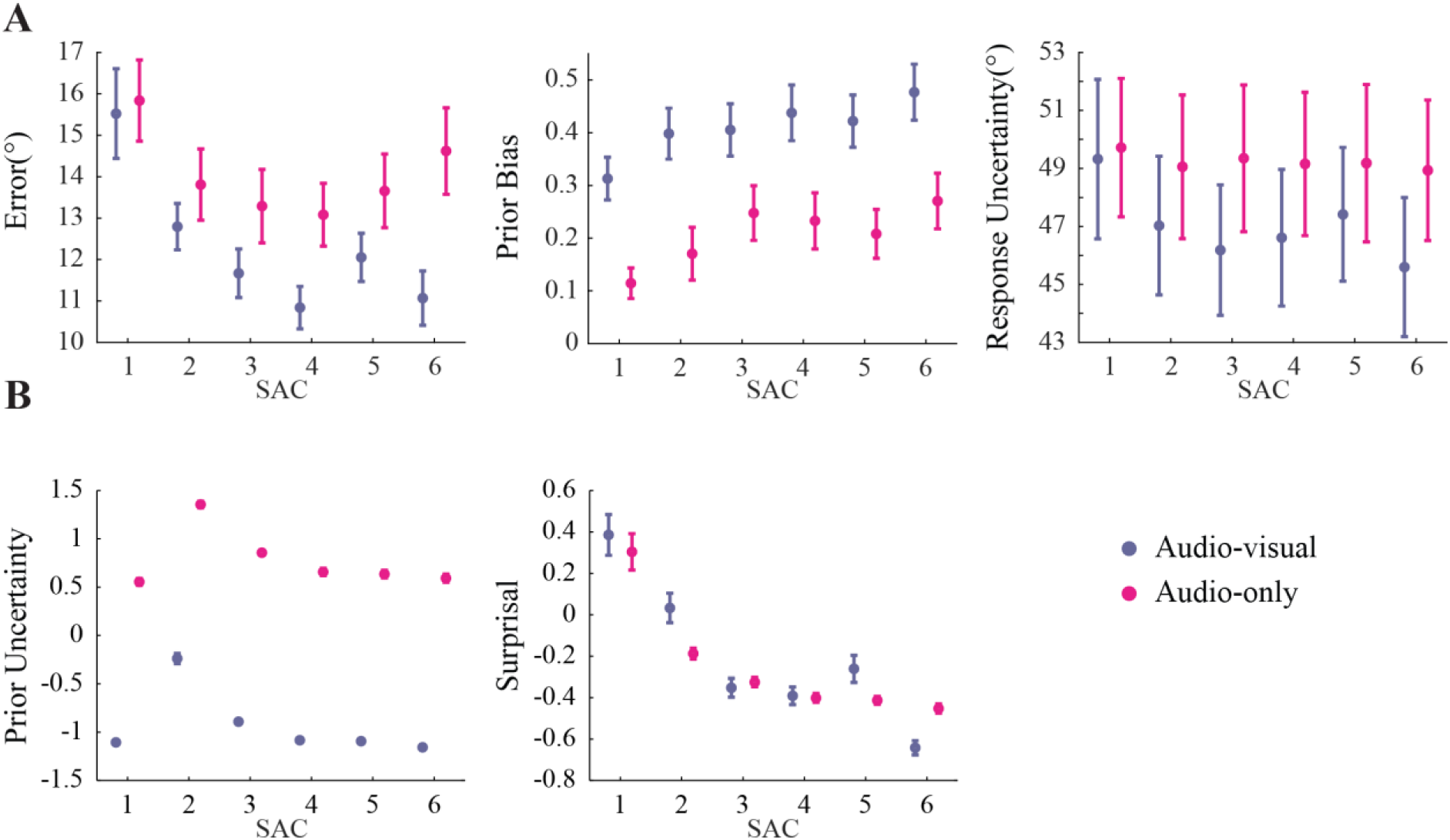
Behavioral and modeling results. **A)** Mean of the behavioral outcomes estimation error, prior bias, and response uncertainty, per sound-after-changepoint (SAC) level, for both audio-visual and auditory conditions. Error bars indicate the standard error of the mean. **B)** Median of the Prior Uncertainty^1^ (PU) and Surprisal (SU) values. Single subjects’ PU and SU values are z-scored. Note, that the SEM is particularly small (mean = .59) for PU in the AV condition. This is due to the fact, that while we fitted the sensory noise individually for each participant for the A condition, we fixed the sensory noise at one degree for the AV condition. Furthermore, we used the same set of trials for all participants, resulting in no variation across the trial sets. Hence, for a particular stimulus, our model predicts the same PU for every participant. The only remaining variation stems from the random sensory noise of the AV stimuli.

**Table 1.** Behavioral results Statistical results obtained using linear mixed-effect modeling.

| Fixed effects coefficients | Estimate | SE | z | Pr(>\|z\|) |
| --- | --- | --- | --- | --- |
| (Intercept) | -1.37 | 0.82 | -1.69 | 0.09 . |
| SAC | 0.2 | 0.17 | 1.17 | 0.24 |
| Modality=AV | 2.56 | 0.89 | 2.88 | <4e-3 ** |
| SAC:Modality=AV | -0.46 | 0.24 | -1.92 | 0.05 |

**B. Response uncertainty**Formula: RU $\sim 1 + \text{SAC} * \text{Modality} * (1 | \text{Subjects})$
| Fixed effects coefficients | Estimate | SE | z | Pr(>\|z\|) |
| --- | --- | --- | --- | --- |
| (Intercept) | 49.52 | 2.44 | 20.29 | <2e-16 *** |
| SAC | -0.09 | 0.14 | -0.63 | 0.53 |
| Modality=AV | -1.06 | 0.76 | -1.4 | 0.16 |
| SAC:Modality=AV | -0.35 | 0.2 | -1.71 | 0.09 |

**C. Prior bias**Formula: Bias $\sim 1 + \text{SAC} * \text{Modality} * (1 | \text{Subjects})$
| Fixed effects coefficients | Estimate | SE | z | Pr(>\|z\|) |
| --- | --- | --- | --- | --- |
| (Intercept) | 0.12 | 0.05 | 2.64 | 8.22e-3 ** |
| SAC | 0.02 | 0.01 | 3.38 | 7.2e-4 *** |
| Modality=AV | 0.2 | 0.04 | 5.23 | 1.65e-07 *** |
| SAC:Modality=AV | <3e-4 | 0.01 | 0.02 | 0.98 |

**D. Prior uncertainty**Formula: PU $\sim 1 + \text{SAC} * \text{Modality} * (1 | \text{Subjects})$
| Fixed effects coefficients | Estimate | SE | z | Pr(>\|z\|) |
| --- | --- | --- | --- | --- |
| (Intercept) | 16.57 | 0.3 | 54.37 | <2e-16 *** |
| SAC | -0.37 | 0.02 | -15.21 | <2e-16 *** |
| Modality=AV | -7.64 | 0.13 | -59.52 | <2e-16 *** |
| SAC:Modality=AV | -0.04 | 0.03 | -1.14 | 0.25 |

**E. Surprisal**Formula: SU $\sim 1 + \text{SAC} * \text{Modality} * (1 | \text{Subjects})$
| Fixed effects coefficients | Estimate | SE | z | Pr(>\|z\|) |
| --- | --- | --- | --- | --- |
| (Intercept) | 5.83 | 0.1 | 57.74 | <2e-16 *** |
| SAC | -0.17 | 0.01 | -13.86 | <2e-16 *** |
| Modality=AV | 0.3 | 0.07 | 4.49 | 7.02e-06 *** |
| SAC:Modality=AV | -0.01 | 0.08 | -0.5 | 0.62 |

**Table 2.** Cluster p-values obtained from regression analysis.

| **A. Time domain regression clusters** |  |  |  |  |  |  |  |
| --- | --- | --- | --- | --- | --- | --- | --- |
|  | **C1** | **C2** | **C3** | **C4** | **C5** | **C6** | **C7** |
| **AV PU** | 0.001 | 0.001 | 0.001 | 0.001 |  |  |  |
| **A PU** | 0.001 | 0.001 | 0.001 | 0.001 | 0.017 | 0.006 |  |
| **AV SU** | 0.012 | 0.005 | 0.001 | 0.001 | 0.007 | 0.013 | 0.011 |
| **A SU** | 0.004 | 0.002 | 0.001 | 0.002 | 0.002 | 0.017 | 0.004 |
| **B. Time-frequency domain lower frequency range (4-30 Hz) regression clusters** |  |  |  |  |  |  |  |
|  | **C1** | **C2** | **C3** | **C4** | **C5** | **C6** | **C7** |
| **AV PU** | 0.001 | 0.001 | 0.001 | 0.002 |  |  |  |
| **A PU** | 0.001 | 0.023 | 0.024 |  |  |  |  |
| **AV SU** | 0.014 | 0.018 | 0.001 | 0.001 | 0.001 | 0.001 | 0.006 |
| **A SU** | 0.001 | 0.001 | 0.001 | 0.001 | 0.013 |  |  |
| **C. Time-frequency domain higher frequency range (40-100 Hz) regression clusters** |  |  |  |  |  |  |  |
|  | **C1** | **C2** | **C3** | **C4** | **C5** | **C6** | **C7** |
| **AV PU** | 0.001 |  |  |  |  |  |  |
| **AV SU** | 0.001 | 0.002 | 0.001 | 0.003 | 0.023 | 0.001 | 0.005 |

**Table 3.** Cluster p-values obtained from condition comparison analysis.

| **A. Time domain regression clusters** |  |  |  |  |  |  |  |
| --- | --- | --- | --- | --- | --- | --- | --- |
|  | **C1** | **C2** | **C3** | **C4** | **C5** | **C6** | **C7** |
| **AV-A PU** | 0.004 | 0.001 | 0.001 | 0.001 | 0.005 | 0.022 |  |
| **AV-A SU** | 0.007 | 0.002 | 0.006 | 0.002 | 0.001 | 0.014 |  |
| **B. Time-frequency domain lower frequency range (4-30 Hz) regression clusters** |  |  |  |  |  |  |  |
|  | **C1** | **C2** | **C3** | **C4** | **C5** | **C6** | **C7** |
| **AV-A PU** | 0.002 | 0.013 |  |  |  |  |  |
| **AV-A SU** | 0.001 | 0.004 | 0.001 | 0.015 |  |  |  |
| **C. Time-frequency domain higher frequency range (40-100 Hz) regression clusters** |  |  |  |  |  |  |  |
|  | **C1** | **C2** | **C3** | **C4** | **C5** | **C6** | **C7** |
| **AV-A SU** | 0.002 | 0.001 | 0.004 |  |  |  |  |

For the latent model variable PU, the LMM analysis revealed a main effect of Modality (*p* < .001). This was due to larger PU values in the A compared to the AV condition, in line with our expectation of a weaker prior in the former. Further, we found a main effect of SAC level (*p* < .001), with the largest PU values at the second sound after a change-point, indicating that change-points led to a transient increase in PU.

For the model variable SU, we found a significant main effect of Modality (*p* < .001), due to overall larger SU values in the AV compared to the A condition. Again, this is likely a consequence of a stronger prior in the former, which leads to stronger surprisal following a change-point. Finally, we observed a main effect of SAC level (*p* < .001), due to surprisal being largest directly after a change-point, and then decreasing with increasing numbers of sounds after the change-point. No interactions were observed for the model variables (*p* > .25).

### EEG regression results

Figure 3 displays the results of the regression (i.e. beta coefficients) between the time-domain EEG data and the model variables PU and SU (see Supplementary Table 2 for the cluster statistics). For both the AV and A condition, there are two time periods in which PU significantly predicts the EEG activity patterns. The first is between around -250 to -100 ms prior to sound onset, in line with activated prior information in expectation of a stimulus. The second period is between 0 and 100 ms following a sound, potentially tracking the comparison between the prior and the sensory stimulus. Further, both periods are comprised of a medio-central- and an occipital cluster of electrodes with opposing beta coefficient values, likely due to the dipolar pattern of an underlying event-related potential.

**Figure 3.**
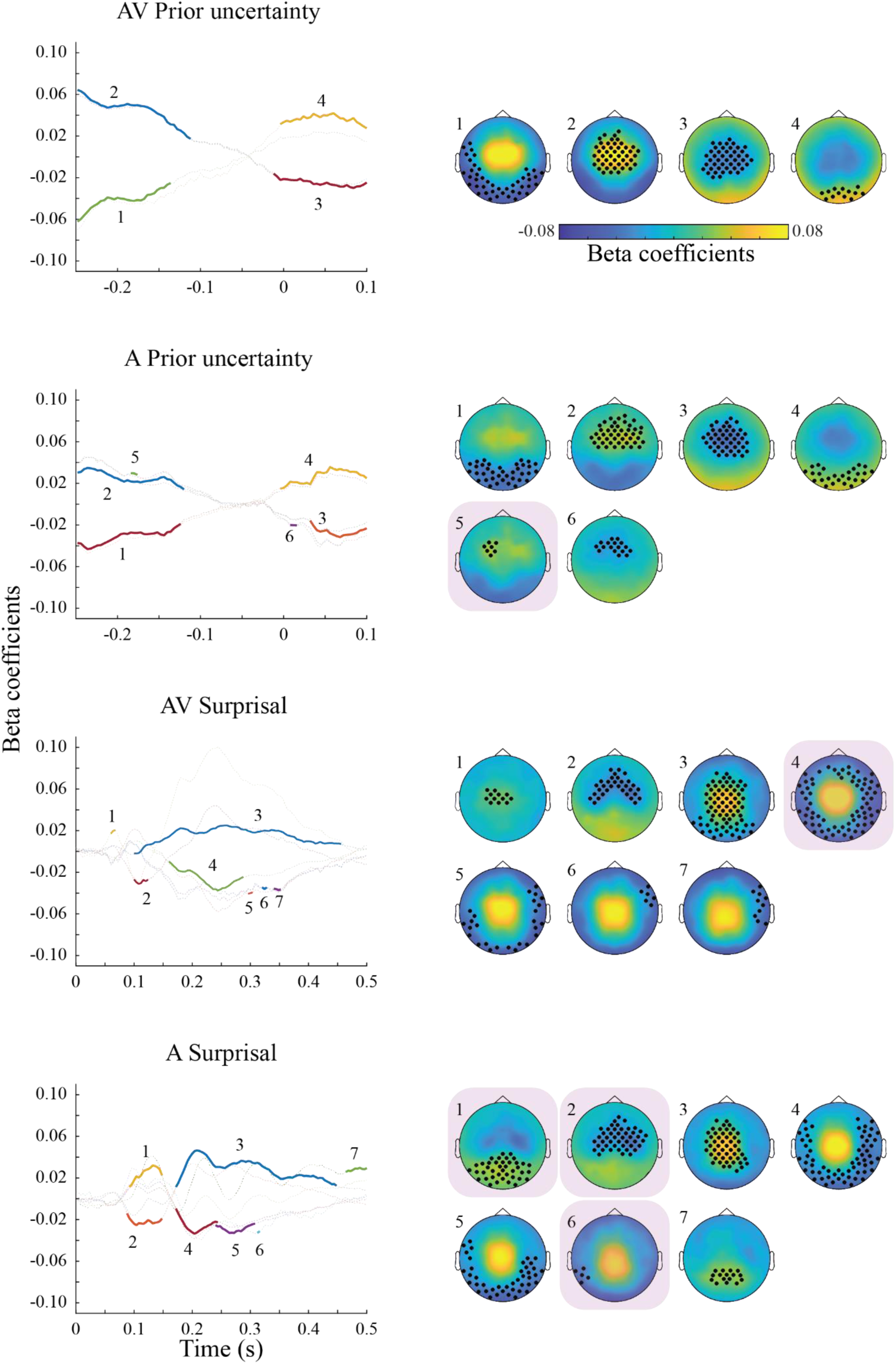
Event-related responses associated with Prior Uncertainty (PU) and Surprisal (SU). Left column: Time-domain representations of clusters (numbered) of regression coefficients between EEG amplitudes and model latent variables, significant at the group-level. Data are shown separately for the audio-visual (AV) and auditory (A) conditions, as well as for PU and SU variables. Each line shows beta coefficients averaged over channels and time points of each significant cluster, obtained via permutation testing. Significant time points are marked in bold colors. Right column: Topographical plots of each significant cluster, as numbered on the left side as. Significant channels are marked with asterisks. Behaviorally relevant clusters (see Figure 5A) are highlighted via a purple square in the background. Small clusters (< 2 datapoints) are not shown, unless they are behaviorally relevant.

For the regression against SU, both AV and A conditions show significantly associated EEG time-domain patterns aggregated between 100 and 500 ms following a sound, in line with a surprisal response to the comparison between prior and sensory input. Again, the topography shows a dipolar pattern, with medio-central and occipital clusters of opposing beta coefficients. In summary, the time domain regression shows large, spatially overlapping, but temporally distinct EEG activity patterns associated with PU and SU.

In the time-frequency domain (Figure 4), regressions revealed that pre-stimulus synchronization in the delta-theta range was positively associated with PU, in both AV and A conditions. Only for the AV condition, PU was negatively associated with synchronization around the stimulus onset in the alpha and beta frequency range. Finally, also in the AV condition, pre-stimulus gamma power (∼75 Hz) was positively associated with PU.

**Figure 4.**
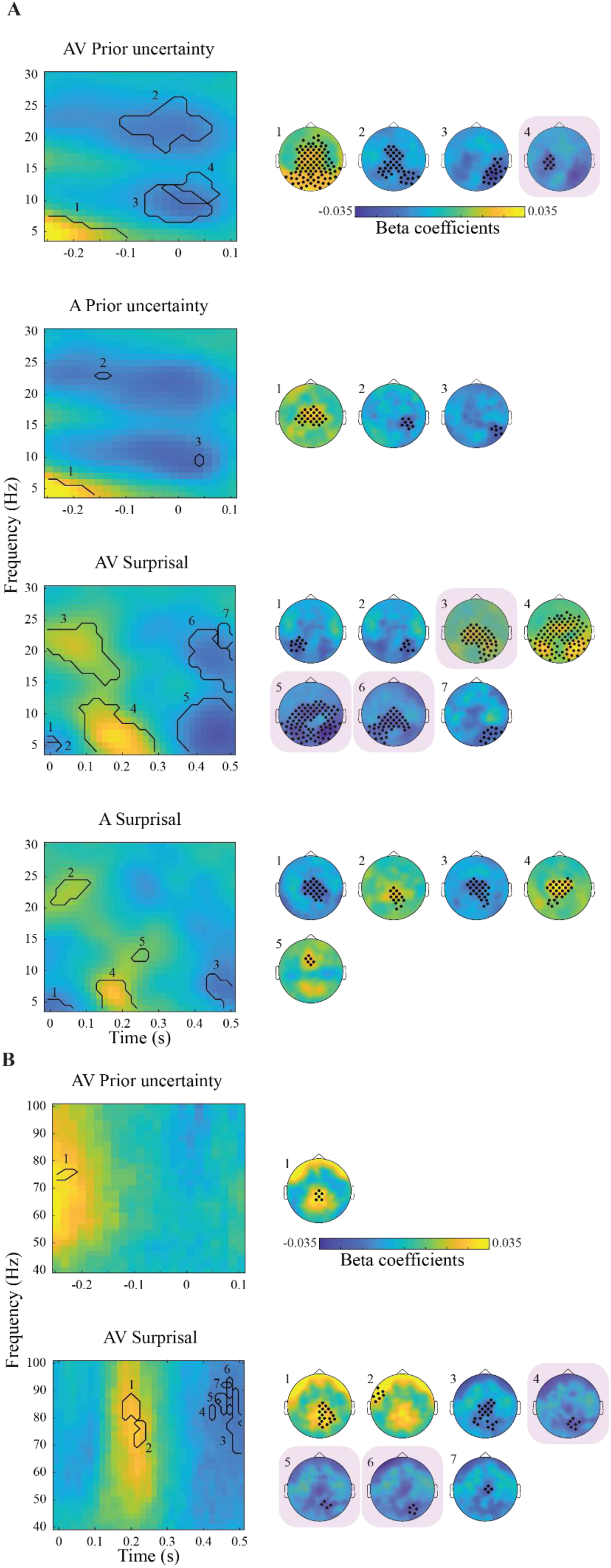
**A)** Time-frequency responses (lower frequency range, 4-30 Hz) associated with Prior Uncertainty (PU) and Surprisal (SU). Left column: Time-frequency representations of regression coefficients between EEG amplitudes and model latent variables, averaged over all significant channels obtained via permutation testing. Significant clusters at the group-level are highlighted and numbered. Data are shown separately for the audio-visual (AV) and auditory (A) conditions, as well as for PU and SU variables. Right column: Topography plots of each significant cluster, as numbered on the left side as. Significant channels of the clusters are marked with asterisks. Behaviorally relevant clusters (see Figure 6B) are highlighted via a purple square in the background. **B)** Same as A, for higher frequency range (i.e. 40-100 Hz). Small clusters (< 2 datapoints) are not shown, unless they are behaviorally relevant.

Several time-frequency activity patters are also significantly linked to SU values. In both the AV and A condition, SU is associated with reduced low frequency activity around the initial onset of the stimulus. Subsequently, however, around 200 ms, the relationship becomes positive for the theta to alpha range, arguably due to enhanced processing of surprising stimuli. Likewise, early post-stimulus activity also correlates positively with beta-band power. During the later time period around 400 – 500 ms following the stimulus, synchronization in the theta-alpha as well as in the beta range (the latter only for AV) is negatively associated with SU. Lastly, gamma-band power (∼70 – 90 Hz) in the AV condition was positively associated with SU at around 200 ms following the stimulus, and negatively associated later between 400 and 500 ms post-stimulus.

As in the time-domain, PU and SU had temporally distinct, but spatially and spectrally overlapping associated EEG activity patterns, suggesting a processing cascade from the representation of the prior to the computation of a prediction error and a subsequent surprise.

### Neural patterns in the AV versus A condition

Overall, regression coefficients in both time domain and time-frequency domain were larger for the AV compared to the A condition, resulting in more and larger significant clusters. In the time domain, comparison of the regression coefficients associated with PU between the two conditions revealed two centrally located positive clusters and four negative fronto-temporal clusters in the pre-stimulus time period (Figure 5). In the time-frequency domain, we found an earlier positive posterior cluster (∼ -250 to -50ms) in theta range and a negative frontal cluster in beta range (∼12-15 Hz) starting around 100 ms prior to the sound onset. No significant clusters were found for PU in the higher frequency range. For SU in the time domain, we found three positive and three negative clusters between ∼150 to 300ms, possibly reflecting larger underlying ERPs in the AV compared to the A condition. In the lower frequency range, one negative fronto-central cluster in beta range (∼12-14 Hz), and two negative clusters and one positive occipital cluster in theta range were found for SU, similarly due to the larger coefficients in AV condition. Finally, in the high frequency range we observed three positive clusters (∼150 to 220 ms) between ∼ 66-90 Hz associated with SU, indicating larger coefficient values in the AV condition.

**Figure 5.**
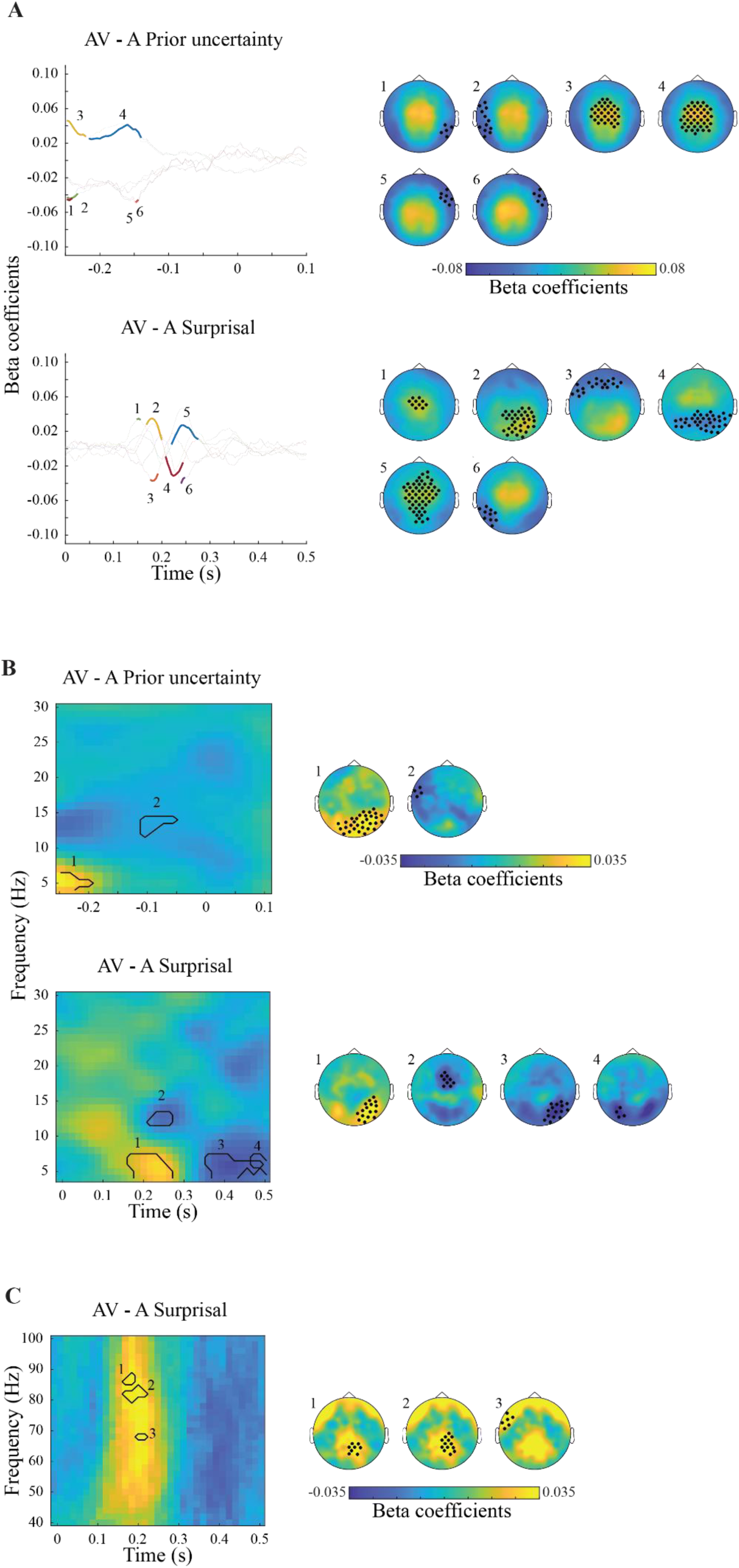
Differences in Prior Uncertainty (PU) and Surprisal (SU) related activity between the audio-visual (AV) and auditory (A) conditions. All data represent the difference of regression coefficients (between the EEG data and the PU and SU latent variables, see Figures 4 and 5) between the AV and A conditions. **A)** Left column: Time-domain representations of regression coefficient differences. Data are shown separately for the PU and SU variables. Each line shows beta coefficient differences averaged over channels and time points of each significant cluster, obtained via permutation testing. Significant time points are marked in bold colors. Right column: Topographical plots of each significant cluster, as numbered on the left side as. Significant channels are marked with asterisks. **B)** Left column: Time-frequency (4 – 30 Hz) domain representations of regression coefficient differences. Data are shown separately for the PU and SU variables. Significant clusters at the group-level are highlighted and numbered. Data are shown separately for the PU and S variables. Right column: Topography plots of each significant cluster, as numbered on the left side as. Significant channels of the clusters are marked with asterisks. **C)** Same as B, for the 40-100 Hz time-frequency domain. Small clusters (< 2 datapoints) are not shown.

### Behavioral relevance of PU and SU related neural activity

After establishing the above distinct PU- and SU-related activity patterns for both the AV and A conditions, we tested which of those patterns were, on a trial-by-trial level, predictive of the behavioral metrics (error, response uncertainty, prior bias; see Figure 6). Importantly, as detailed in the Methods section, we did so by regressing the behavioral variables against the EEG data around the final probe sound of each trial. Thus, these data are independent of the data used for the regression against the model variables reported above, which are taken from each sound of the sequence *except* the final probe sound.

**Figure 6.**
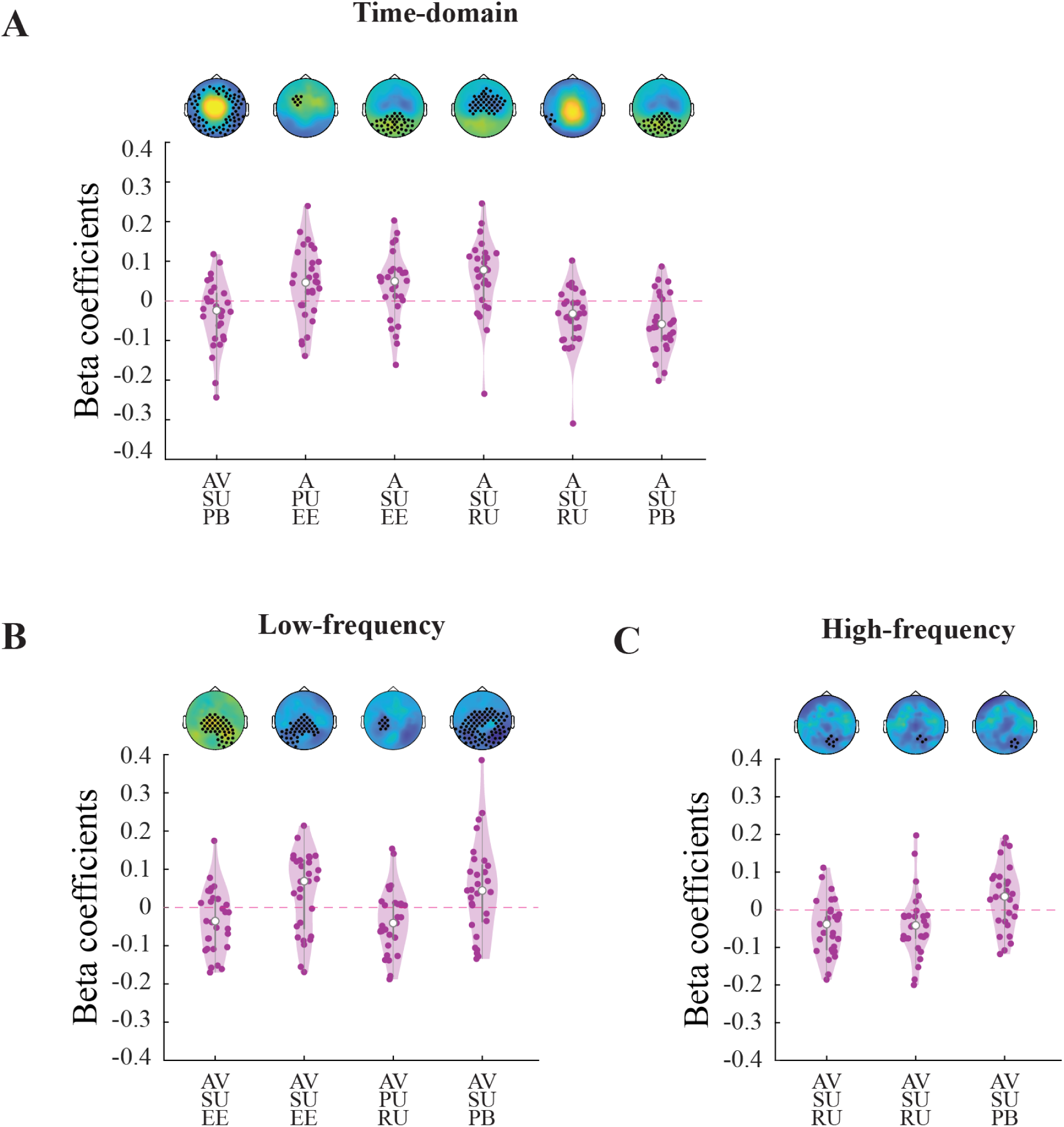
Model latent variable components predict behavior. A) Results of the regression analysis between time-domain EEG data averaged over the dimensions of significant regression clusters (see Figure 4) and the behavioral estimation error (EE), response uncertainty (RU), and prior bias (PB) variables. Each circle in the violin plots represents single subject beta coefficients. Grey bars represent the median, interquartile range and 1.5 times the interquartile range. Respective topoplots of the significant clusters are shown on top of the violin plots (same as the highlighted clusters in Figures 4). B) Same as A), for the 4-30 Hz time-frequency domain clusters (see Figure 5). C) Same as B), for the 40-100 Hz time-frequency domain clusters (see Figure 6).

The participants’ estimation error in the A condition was significantly predicted by the activity of a frontal, pre-stimulus time-domain cluster positively associated with PU, as well as an occipital post-stimulus cluster positively associated with SU.

In the AV condition, estimation error was predicted by an early (0-150ms) posterior beta-band cluster positively associated with SU and a later (400 – 500ms) posterior beta-band cluster negatively associated with SU.

Prior bias in the A condition was predicted by a posterior time-domain cluster (100 – 200ms) positively associated with SU. In the AV condition, prior bias was predicted by a large time-domain cluster (150 – 300ms) negatively associated with SU. As can be seen from Figure 4, topographies and time-courses are similar for both AV and A SU time-courses, yet behavioral correlation with prior bias became significant for two different, consecutive time periods for the two conditions, with different ERP polarities. Thus, although their association to SU is oppositional, these clusters are likely part of the same process, which switches EEG amplitude polarity at around 180ms. In the AV condition, prior bias was further predicted by two time-frequency activity clusters. First, a large posterior cluster in the theta-alpha range (350 – 500ms), in which a desynchronization was associated with more SU and eventually, less prior bias. Second, a small left-posterior cluster in the gamma range (∼470ms), in which a desynchronization was likewise associated with more SU and eventually, less prior bias. Finally, the participants’ response uncertainty ratings in the A condition were predicted by two clusters in the time-domain. First, an early (∼75 to 150ms) large central cluster and second, a small later (∼320ms) posterior cluster, in both of which negative ERP amplitudes were associated with more surprisal and more response uncertainty. In the AV condition, response uncertainty was associated with three clusters in the time-frequency domain: A central-left cluster in the alpha-beta range (∼ -50 – 100ms), in which a desynchronization was associated with more PU as well as more response uncertainty; and two occipital clusters in the gamma-range (∼425 – 450ms), in which a desynchronization was associated with more SU as well as less response uncertainty.

## Discussion

In the present study, we investigated behavioral and neural evidence for Bayesian inference during auditory spatial localization in dynamic environments.

We report three main findings. First, our results show that participants continuously integrated prior knowledge into their estimations, subject to dynamic changes in a volatile environment. Second, these patterns of results were similar but amplified by the presence of additional visual location priors. Third, we observed distinct EEG activity patterns associated with Prior Uncertainty (PU) and Surprisal (SU), which are in line with previous studies using visual and/or temporal or pitch-related tasks. Importantly, these EEG activity patterns were predictive of the error, response uncertainty, as well as the prior bias of our participants’ behavioral responses on a trial-by-trial basis.

### Dynamic environmental changes impact behavioral and Bayesian model data

In our experiment, we used random change-points and experimental noise to simulate a noisy dynamic environment with momentary changes to the reliability of prior information and sensory evidence. Data from both experimental conditions showed stronger prior bias with increasing SAC levels (i.e. accumulated sensory evidence). A similar effect was present for PU, however with a peak at SAC level 2, as expected given the change-point at the previous sound. Together, these results indicate that participants rely more on prior information, as sensory evidence is accumulated during a period of stable environmental statistics and the prior uncertainty decreases.

At the same time, estimation error and SU were largest at SAC level 1 (i.e. immediately following a change-point). The large SU is indicative of a now irrelevant prior, which cannot be used to improve the localization performance anymore, indicating a reason to update the internal model. This was again similarly the case for both the AV and the A condition, suggesting a comparable mechanism for visual and auditory spatial inference (see below for a detailed discussion of the condition differences).

Our findings thus corroborate and extend the results by Krishnamurthy et al.^21^ who reported similar effects following AV stimulation, by demonstrating Bayesian-like inference in unimodal auditory settings and showing visual stimuli likely improved the sound localization performance by providing a more precise prior distribution.

### Prior Uncertainty is coded around stimulus onset, and associated with localization performance and response uncertainty

In the time-domain, the pre-stimulus neural patterns associated with PU likely reflect late ERP responses to the previous stimulus, with their topography and latency suggesting a P3-like component. The P3 is well documented to scale with surprisal and internal model updating^8,18–20^, which fits well with our observed positive association with subsequently increased PU. In the A condition, activity in this time-period was predictive of the behavioral performance, indicating that larger P3-like ERP responses (reflecting larger surprisal due to a prediction error) to the previous stimulus were associated with larger estimation errors in the current trial. In the post-stimulus period, the significant clusters span the time-range of early sensory processing including the P1 component. These scale negatively with PU, thus, the stronger the prior, the larger the resulting sensory evoked ERP. Importantly, this early PU regression pattern is still independent of the prediction error and any resulting surprisal, both of which would be expected to appear later in the time period of a P3 ERP component and scale positively with the ERP response.

The above discussed pre-stimulus ERP likely underlies the theta-band time-frequency domain result, with which it shares a similar topography and time window. Additionally, low frequency oscillations have been associated with temporal expectancy of important upcoming stimuli^40^. Considering the regular SOAs in our study (i.e. always 500 ms), their oscillatory power in our results scale positively with PU, in line with stronger expectation of an upcoming sensory event in times of weak prior information.

Further, in line with numerous previous studies^14,15^, we found prior uncertainty reflected in adjacent alpha- and beta-band activity patterns. Generally, the expectancy for an upcoming task-relevant stimulus is known to cause a decrease in alpha- and beta power, especially for regular inter-stimulus intervals^40^. Thus, as PU increases, we would expect participants to increase the weight of sensory evidence rather than the prior, with the observed decrease in alpha/beta power reflecting attentional preparation of the upcoming stimulus. Moreover, a post-stimulus alpha/beta range cluster was both negatively associated with PU and response uncertainty, suggesting a common neural pattern underlying the prior and the resulting subjective response uncertainty.

Interestingly, previous literature suggests a possible dissociation of neural patterns between predictions and their precision, reflected in beta and alpha range oscillations, respectively^13–15^. Presently, however, we found neural patterns associated with PU in multiple frequency bands in pre-stimulus as well as post-stimulus period. A possible reason for this discrepancy is the applied analysis pipeline. For instance, Sedley et al.^15^ partialized out the correlation between predictor variables prior to their regression analysis, which might have canceled out correlation in other frequency bands. Indeed, upon performing the same analysis without the partialization, they similarly observed an effect for prior precision in delta/theta, alpha and beta/gamma-bands.

Finally, we found a positive gamma cluster in the pre-stimulus time period for PU. Considering the short SOAs in our study (500ms), this association might be related to the previous stimulus processing reflecting facilitation of sensory processing and increased weighting of the sensory likelihood in response to larger prior uncertainty and reduced prior reliability.

However, the observed time-window (∼250ms following the previous stimulus), the broadband nature, as well as its presence predominantly in the AV condition, indicate that the gamma-response might be affected by microsaccadic muscular activity^41^. Although we diligently removed eye-related independent components, which can help to clear microsaccadic artifacts from EEG data^42^, we are cautious to interpret the gamma response in this time-period as genuinely brain-related.

### Surprisal-related EEG activity evolves post-stimulus and is linked to behavioral error, confidence and prior bias

As surprisal (SU) represents the deviation of sensory evidence from the prior, we consequently expected it to be represented in post-stimulus neural activity patterns. In the past, EEG studies repeatedly found P3-like responses in response to surprising events 8,18-20, whose amplitude scales with the amount of surprisal and subsequent internal updating, and which have been interpreted as the supporting evidence for the Bayesian brain hypothesis^43^. In line with these results, SU in our data was associated with neural activity in the time range of the P3 ERP component, in both AV and A conditions. Importantly, this adds to the previous literature by showing SU related responses in auditory spatial inference tasks, despite the auditory systems inferiority regarding localization, and thus demonstrates the ubiquity of the associated P3-related mechanism.

In both the A and AV condition, larger amplitudes in early SU-related time-domain clusters (∼100-300 ms) were predictive of less prior bias in behavioral responses, indicating less use of prior information following stronger neural correlates of surprisal. In addition, subjective response uncertainty was likewise significantly positively associated with activity in two SU-related time-domain clusters, although only in the A condition. As to be expected, stronger activation of these patterns subsequently led to larger response uncertainty, on a trial-by-trial basis.

Surprisal has further been related to gamma-band oscillations, possibly reflecting the processing of new sensory input^14–16^. Indeed, in the AV condition we found several clusters in the 70 – 90 Hz frequency range, associated with SU. While two of those falls into the time-range of 200 – 300ms and are potentially driven by microsaccadic activity (as discussed above), three of the later ones between 400 and 500 ms are particularly interesting, as their activity correlates with subsequent response uncertainty and bias. More specifically, larger gamma-band power in these clusters is associated with less surprisal and in subsequent behavior, larger prior bias and smaller response uncertainty.

In addition to what previous studies have reported, our data also revealed SU to be reflected in lower frequency activity. The activity patterns across space, time and frequency were very similar between the AV and A conditions, with overall more pronounced power changes in the former. Starting from about 400ms after stimulus onset, alpha/beta-band power is negatively correlated with SU. High levels of surprisal indicate change-points which render the current prior irrelevant. Thus, with large SU levels alpha/beta range power decreases in order to facilitate processing of the upcoming stimulus. In line with this interpretation, two clusters in the AV condition’s alpha/beta range were predictive of the subsequent behavioral estimation error and prior bias, respectively.

Several other clusters associated with surprisal were also predictive of behavioral outcomes. Interestingly, activity in an early beta-band cluster in the AV condition scaled positively with SU and subsequent localization accuracy (i.e. negatively with estimation error). The fact that stronger neural correlates of SU would be associated with better performance is suggestive of an attention effect, in that the appearance of a salient change-point leads to surprisal, but concurrently boosts attention and improves task performance.

The fact that we did not find similar behaviorally relevant clusters in the A condition might be due to the lower signal-to-noise ratio as well as the lower stimulus intensity following an A-only compared to a combined auditory and visual stimulus^44^.

### Behavioral and neural tokens of Bayesian inference are more pronounced following AV compared to A stimulation

Overall, the location estimation performance was better for the AV compared to the A condition, despite the identical probe sounds (both audio-only), in line with previous studies^22,23^. In part, this result likely reflects differences in the intrinsic sensory reliability of visual and auditory spatial information^28,29^. Additionally, A condition was particularly challenging due to the auditory-to-visual response mapping. Despite this difference, both conditions showed response bias towards the prior stimulus locations, indicating participants indeed kept track of the previous locations.

However, the higher sensory noise in the A-only condition led to larger PU and therefore less prior bias during sound localization. In the AV condition, formation of the prior information relies on both senses and as a result, participants relied on their audio-visual prior more than the noisy audio-only sensory evidence provided by the probe sound. Similarly, a more precise prior in the AV condition potentially led to larger SU values for the AV compared to the A condition.

Overall, our behavioral results indicate similar mechanism for integration of prior and sensory likelihood following unimodal A and bimodal AV stimulation, that complies with Bayesian inference in a modality and task independent manner.

Likewise, the comparison of the neural activity associated with PU and SU between the conditions revealed largely similar patterns. In both time- and time-frequency domain, the observed differences were most likely due to the larger and more extensive regression coefficient patterns for the AV compared to the A condition, due to stronger sensory activations and consequently a higher signal-to-noise ratio in the former.

For PU, the comparison of the two conditions in the time domain revealed clusters in the pre-stimulus time period due to a more pronounced PU representation in the AV compared to the A condition. In the time-frequency domain, the two conditions have spatially more distinct PU patterns in theta range, reflected by a significant occipital cluster in comparison analysis, that likely stems from the additional visual prior representation in the AV condition.

Comparison analysis revealed similar results for SU, mainly reflecting differences in strength of the correlations between the two conditions. Additionally, these differences might partly also be driven by SU itself, which showed larger variation in the AV compared to the A condition. In the time domain, the differences are in the time range of ∼200-300 ms due to more pronounced regression patterns in the AV compared to the A condition. Similarly, in lower frequency range, the differences between the two conditions are mainly in the theta range and limited to occipital electrodes, possibly due to sensory specific processing of the visual stimulus in the AV condition. Finally, the difference in the gamma range clusters fall into the period of potential microsaccadic activity, and thus are likely driven by larger artifacts in the AV condition due to the occurrence of an additional visual stimulus.

Taken together, these results suggest that auditory spatial localization in both AV and A conditions is based on highly similar Bayesian inference mechanisms, which in the former are additionally informed by the visual stimulus. Importantly, our data thus provide novel evidence for auditory spatial localization according to Bayesian inference, while suggesting that the results following audio-visual priors previously reported by Krishnamurthy et al.^21^ are similarly applicable to true auditory-only settings.

## Conclusions

Perception commonly relies on both sensory input and prior information. In the present study, we provide novel evidence that behavioral and neural responses during unimodal auditory localization indeed conform with Bayesian inference principles. We demonstrate the impact of dynamic changes in the environment on the weighting of prior knowledge and current sensory evidence, and show that the resulting behavioral performance, model metrics and neural patterns associated with PU and SU are in line with findings from other domains. Moreover, these patterns are intensified yet structurally similar following additional visual priors.

Taken together, despite the auditory system’s inferiority regarding spatial localization, our data suggest that it employs similar mechanisms as previously observed in visual processing or more domain appropriate-tasks such as auditory pitch and temporal discrimination, supporting Bayesian inference as a general principle in human perceptual decision making.

## Acknowledgments

This research was supported by an Austrian Science Fund (FWF) Young Independent Researchers Group (Grant-DOI: 10.55776/ZK66) to Michelle Spierings, Ulrich Pomper, and Robert Baumgartner.

## Author Contributions

**Burcu Bayram** Data curation; formal analysis; visualization; writing-original draft

**David Meijer** Data curation; formal analysis; writing-original draft

**Roberto Barumerli** Formal analysis; writing-review and editing

**Michelle Spierings** Conceptualization; funding acquisition; project administration; resources; writing-review and editing

**Robert Baumgartner** Conceptualization; funding acquisition; project administration; resources; writing-review and editing

**Ulrich Pomper** Conceptualization; funding acquisition; project administration; resources; supervision; writing-original draft

## Data Availability Statement

Data are available upon request.

