## Supplement 1 for "Bayesian Prior Uncertainty and Surprisal Elicit Distinct Neural Patterns During Sound Localization in Dynamic Environments"

**Bayesian model**

To retrieve response predictions for each trial and estimates of the internal, latent variables of prior uncertainty and surprisal for each sound, we fitted a Bayesian inference model to the spatial estimation responses of each participant. In this supplementary section, we successively describe the computational model and the fitting procedure.

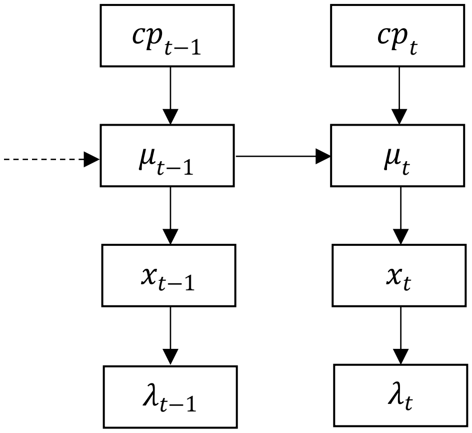


$$\begin{cases} cp_t = 1 & if\ t = 1 \\ cp_t \sim B(1, H) & if\ t > 1 \end{cases}$$

$$\begin{cases} \mu_t = \mu_{t-1} & if\ cp_t = 0 \\ \mu_t \sim U(\mu_t; \alpha, \beta) & if\ cp_t = 1 \end{cases}$$

$$x_t \sim N(x_t; \mu_t, \sigma^2{}_{exp})$$

$$\lambda_t \sim \gamma_\lambda N\left(\lambda_t; x_t, \left(\sigma_{sens,0°} + \gamma_\sigma |x_t|\right)^2\right)$$

**Figure S1**. Generative model. The graphic illustration (left) and accompanying formulas (right) describe the generative process of an observer's sensory signals. See text below for details.

<u>Generation of sensory signals</u>

The stimuli sequence of every trial was constructed using a pseudorandom algorithm that is displayed in supplementary Figure S1. We sampled the first generative mean for each trial, $\mu_{t=1}$, from a bounded uniform distribution between by $\alpha = -60°$ and $\beta = +60°$. At all later times, $t > 1$, a random binomial draw determined whether a changepoint occurred ($cp_t = 1$). The probability of a changepoint was given by hazard rate H, which was constant (1/6) for all stimuli (after the first). If a changepoint occurred, then the generative mean $\mu_t$ was resampled from the bounded uniform distribution $U(\mu_t; \alpha, \beta)$. Otherwise, $\mu_t$ was equal to the previous generative mean, $\mu_{t-1}$. Stimulus locations $x_t$ were sampled from a normal distribution, $N(x_t; \mu_t, \sigma^2{}_{exp})$, that was centred on the generative mean, with standard deviation given by a fixed amount of experimental noise $\sigma_{exp} = 10°$.

Participants only have access to noise-corrupted internal sensory estimates $\lambda_t$. We assume normally distributed sensory noise with standard deviation $\sigma_{sens,x_t}$, which may linearly increase with the absolute stimulus location: $\sigma_{sens,x_t} = \sigma_{sens,0°} + \gamma_\sigma |x_t|$, where $\gamma_\sigma \geq 0$. We further allow for possible biases of the internal estimates by multiplication with a gain factor, $\gamma_\lambda$, such that $\lambda_t$ is sampled from $\gamma_\lambda N(\lambda_t; x_t, \sigma^2{}_{sens,x_t})$. Sensory biases can be directed towards ($0 \leq \gamma_\lambda \leq 1$) or away from ($\gamma_\lambda > 1$) the midline.

Bayesian inference about last sound location $x_t$

An observer's task is to infer the true stimulus location $x_t$ at the end of a sequence. The simplest estimate of $x_t$ is directly given by $\lambda_t$. However, this internal estimate was corrupted by random noise and may be biased (if $\gamma_\lambda \neq 1$), so it is unreliable to some extent. Bayesian observers take such uncertainty into account and construct a likelihood function over all possible values for $x_t$ by inverting the generative function. Here, we assume that observers are able to estimate the amount of random noise for a particular stimulus ($\sigma^2{}_{sens,x_t}$), but that they remain unaware of any systematic bias (i.e. they assume that $\gamma_\lambda = 1$). A useful analogy is that of an observer seeing a blurry image in the mirror of a steamy bathroom, but being unaware of the fact that the mirror is also slightly tilted. Under these assumptions, the likelihood function about stimulus location $x_t$ can be represented as a normal distribution:

$$p(\lambda_t|x_t) = N(x_t; \lambda_t, \sigma^2{}_{sens,x_t}) \quad \text{(Eq. 1)}$$

The uncertainty about the true stimulus location, based only on the latest sensory evidence $\lambda_t$, can be defined as the standard deviation of the likelihood function: $\sigma_{sens,x_t}$. Bayesian theory postulates that an observer can minimize this uncertainty, and thus the expected error, by using all available sensory information ($\lambda_{1:t}$) and their knowledge of the generative process. This is because, depending on whether a change-point has taken place ($cp_t$), preceding observations carry information about the current generative mean ($\mu_t$), and thus also about the location of the last stimulus because $p(x_t|\mu_t) = N(x_t; \mu_t, \sigma^2{}_{exp})$. Consequently, if one is able to obtain a best estimate of the location of the generative mean, $\hat{\mu}_t = E(p(\mu_t|\lambda_{1:t}))$, then it can be combined with the latest internal estimate $\lambda_t$ by means of a precision-weighted average to obtain a best estimate of the last stimulus location:

$$\hat{x}_t = \frac{\sigma^2{}_{exp}}{\sigma^2{}_{exp}+\sigma^2{}_{sens,x_t}}\lambda_t + \frac{\sigma^2{}_{sens,x_t}}{\sigma^2{}_{exp}+\sigma^2{}_{sens,x_t}}\hat{\mu}_t \quad \text{(Eq. 2)}$$

This best estimate of the last sound location is equal to the mean of the posterior distribution $\hat{x}_t = E(p(x_t|\lambda_{1:\mathrm{t}}))$. To see the benefit of the Bayesian inference scheme over the method of simply selecting the latest observation as one's best estimate of $x_t$, we also give the variance of the posterior distribution $p(x_t|\lambda_{1:\mathrm{t}})$:

$$\sigma^2{}_{x_t} = \frac{\sigma^2{}_{\mu_t}\sigma^4{}_{sens,x_t}}{(\sigma^2{}_{exp}+\sigma^2{}_{sens,x_t})^2} + \frac{\sigma^2{}_{exp}\sigma^2{}_{sens,x_t}}{\sigma^2{}_{exp}+\sigma^2{}_{sens,x_t}} \quad \text{(Eq. 3)}$$

$$= \sigma^2{}_{sens,x_t}\left(\frac{\sigma^2{}_{\mu_t}}{\sigma^2{}_{exp}+\sigma^2{}_{sens,x_t}}\frac{\sigma^2{}_{sens,x_t}}{\sigma^2{}_{exp}+\sigma^2{}_{sens,x_t}} + \frac{\sigma^2{}_{exp}}{\sigma^2{}_{exp}+\sigma^2{}_{sens,x_t}}\right)$$

Here, $\sigma^2{}_{\mu_t}$ is the variance of the posterior distribution about the generative mean, $p(\mu_t|\lambda_{1:\mathrm{t}})$. Equation 3 shows that the uncertainty about $x_t$ (i.e. expected root mean squared error) will be smaller than $\sigma_{sens,x_t}$ as long as $\sigma^2{}_{\mu_t}$ is smaller than $\sigma^2{}_{exp} + \sigma^2{}_{sens,x_t}$. This is usually the case because the variance of the likelihood function about $\mu_t$, i.e. from a single observation $\lambda_t$, is already equal to ${\sigma^2}_{exp} + {\sigma^2}_{sens,x_t}$:

$$p(\lambda_t|\mu_t) = N(\mu_t; \lambda_t, {\sigma^2}_{exp} + {\sigma^2}_{sens,x_t}) \qquad \text{(Eq. 4)}$$

With multiple observations, evidence about the generative mean $\mu_t$ can be integrated to decrease the uncertainty about $\mu_t$, and thus also decrease the expected error about the last stimulus location.

Tracking generative mean $\mu_t$

In the above, we have seen that it is advantageous for task performance to obtain a precise estimate of the generative mean $\mu_t$. Hence, a Bayesian observer updates the posterior distribution $p(\mu_t|\lambda_{1:t})$ iteratively after every new observation $\lambda_t$. Then at the end of a sequence, when prompted to give a response, the observer computes its best estimate for the generative mean $\hat{\mu}_t$, and subsequently its best estimate for the last sound location $\hat{x}_t$ according to equation 2.

The difficulty in tracking the generative mean is that for any new observation (after the first) there are two options: either (1) there was a change-point and the latest observation thus stems from a new generative mean, rendering the previously collected evidence about $\mu_t$ irrelevant; or (2) there was no change-point and the latest observation should thus be integrated with preceding evidence to improve precision. A Bayesian observer does not choose deterministically between these two options. Instead, it computes a posterior distribution for both and then merges them probabilistically. Hence, the fully Bayesian solution to the change-point problem creates mixture distributions, which are memory intensive and computationally intractable[1]. Instead, we will follow the reduced Bayesian observer approach that was introduced by Nassar et al.[2,3]. This method summarizes the posterior mixture distributions into a single normal distribution at every timestep.

The first option is that of a change-point ($cp_t = 1$). According to the generative model, the conditional prior should be the uniform distribution $U(\mu_t; \alpha, \beta)$. However, we instead approximate that conditional prior as a normal distribution with the same mean and variance as the uniform distribution:

$$p(\mu_t | cp_t = 1) = N\left(\mu_t;\ \frac{\alpha+\beta}{2}, \frac{(\beta-\alpha)^2}{12}\right) \qquad \text{(Eq. 5)}$$

The choice for a normal distribution simplifies the iterative algorithm for Bayesian belief updating about the generative mean $\mu_t$. Since the likelihood function is also normally distributed (Eq. 4), it ensures that the conditional posterior also is a normal distribution with the following mean and variance (double bar on top signifies that these are posterior parameters):

$$\bar{\bar{\mu}}_{cp_t=1} = w_{cp_t=1}\ \frac{A+B}{2} + \left(1 - w_{cp_t=1}\right)\lambda_t \qquad \text{(Eq. 6)}$$

$$\bar{\bar{\sigma}}^2{}_{cp_t=1} = w_{cp_t=1}\ \frac{(B-A)^2}{12} \qquad \text{(Eq. 7)}$$

where the weight for prior versus likelihood is defined by its relative precision:

$$w_{cp_t=1} = \frac{\sigma^2{}_{exp} + \sigma^2{}_{sens,x_t}}{\frac{(\beta-\alpha)^2}{12} + \sigma^2{}_{exp} + \sigma^2{}_{sens,x_t}} \qquad \text{(Eq. 8)}$$

The second option is that of no-change-point ($cp_t = 0$). In this case, the conditional prior distribution is given by the posterior distribution of the previous timestep. In the reduced Bayesian observer model[2,3], this prior is a normal distribution.

$$p\left(\mu_t \middle| \lambda_{1:(t-1)}, cp_t = 0\right) = N(\bar{\bar{\mu}}_{t-1}, \bar{\bar{\sigma}}^2{}_{t-1}) \qquad \text{(Eq. 9)}$$

So, the posterior distribution conditional on no-change-point is also normally distributed and its parameters are computed via precision-weighted integration of prior and likelihood:

$$\bar{\bar{\mu}}_{cp_t=0} = w_{cp_t=0}\ \bar{\bar{\mu}}_{t-1} + \left(1 - w_{cp_t=0}\right)\lambda_t \qquad \text{(Eq. 10)}$$

$$\bar{\bar{\sigma}}^2{}_{cp_t=0} = w_{cp_t=0}\,\bar{\bar{\sigma}}^2{}_{t-1} \quad \text{(Eq. 11)}$$

where:

$$w_{cp_t=0} = \frac{\sigma^2{}_{exp}+\sigma^2{}_{sens,x_t}}{\bar{\bar{\sigma}}^2{}_{t-1}+\sigma^2{}_{exp}+\sigma^2{}_{sens,x_t}} \quad \text{(Eq. 12)}$$

These two conditional posteriors will be weighted by their probability. To do so, a Bayesian observer computes the posterior probability of a change-point according to Bayes' rule:

$$\Omega_t = p(cp_t = 1|\lambda_{1:t}) = \frac{p(\lambda_t|cp_t = 1)H}{p(\lambda_t|\lambda_{1:(t-1)}, cp_t = 0)(1-H)+p(\lambda_t|cp_t = 1)H} \quad \text{(Eq. 13)}$$

where the priors are defined by the change-point hazard rate $H$, and the likelihoods are given by the probability density of the following normal distributions, evaluated at internal estimate $\lambda_t$:

$$p(\lambda_t|cp_t = 1) = N\left(\lambda_t; \frac{A+B}{2}, \frac{(\beta-\alpha)^2}{12} + \sigma^2{}_{exp} + \sigma^2{}_{sens,x_t}\right) \quad \text{(Eq. 14)}$$

$$p(\lambda_t|cp_t = 0, \lambda_{1:(t-1)}) = N(\lambda_t; \bar{\bar{\mu}}_{t-1}, \bar{\bar{\sigma}}^2{}_{t-1} + \sigma^2{}_{exp} + \sigma^2{}_{sens,x_t}) \quad \text{(Eq. 15)}$$

Finally, the weighted mixture distribution of the two conditional posteriors is then summarized into a single normal distribution with parameters that are equal to the mean and variance of the mixture[3]:

$$\bar{\bar{\mu}}_t = \Omega_t\bar{\bar{\mu}}_{cp_t=1} + (1-\Omega_t)\bar{\bar{\mu}}_{cp_t=0} \quad \text{(Eq. 16)}$$

$$\bar{\bar{\sigma}}^2{}_t = \Omega_t\bar{\bar{\sigma}}^2{}_{cp_t=1} + (1-\Omega_t)\bar{\bar{\sigma}}^2{}_{cp_t=0} + \Omega_t(1-\Omega_t)\left(\bar{\bar{\mu}}_{cp_t=0} - \bar{\bar{\mu}}_{cp_t=1}\right)^2 \quad \text{(Eq. 17)}$$

So, the posterior distribution about the generative mean, $p(\mu_t|\lambda_{1:t})$, is represented by a normal distribution with mean $\bar{\bar{\mu}}_t$ and variance $\bar{\bar{\sigma}}^2{}_t$. If no response is required, then this posterior will be the prior distribution for the next timestep, conditional on no-change-point (Eq. 9). However, if a response is required, then we need to compute a best estimate of the generative mean, $\hat{\mu}_t$, such that it can be used to compute the intended response location $\hat{x}_t$ (according to equation 2).

Compute best estimate of the last generative mean: $\hat{\mu}_t$

So far, the reduced Bayesian observer model has largely ignored the spatial boundaries (α, β) for the generative mean; n.b. $\mu_t$ is sampled from $U(\mu_t; \alpha, \beta)$. This enables the above simplified (i.e. fast) belief updating algorithm for on-the-fly tracking of the generative mean, independent of the sensory modality of the stimuli. However, the behavioral data demonstrated that our participants did not ignore these spatial limits when they made their responses as there were far fewer responses outside of them. We assume that they learned to use these boundaries as spatial anchors when they map their estimate to the semicircular response arc that appeared on screen when they were prompted for a response (i.e. when time allowed for more elaborate computations). To account for the spatial boundaries of the generative mean we split the posterior normal distribution into three probability masses: one in the middle and one on either side of the spatial boundaries.

$$p_1 = \int_{\mu_t=\alpha}^{\beta} N(\mu_t;\ \bar{\bar{\mu}}_t, \bar{\bar{\sigma}}^2{}_t) d\mu_t \qquad \text{(Eq. 18)}$$

$$p_2 = \int_{\mu_t=-\infty}^{\alpha} N(\mu_t;\ \bar{\bar{\mu}}_t, \bar{\bar{\sigma}}^2{}_t) d\mu_t \qquad \text{(Eq. 19)}$$

$$p_3 = \int_{\mu_t=\beta}^{\infty} N(\mu_t;\ \bar{\bar{\mu}}_t, \bar{\bar{\sigma}}^2{}_t) d\mu_t \qquad \text{(Eq. 20)}$$

The probability masses from outside of the spatial limits ($p_2$ and $p_3$) are assigned to the location of the spatial boundaries themselves. The three parts together thus form a weighted mixture distribution of one doubly truncated normal distribution and two Dirac delta functions at its edges. The best estimate for the generative mean is then computed as the mean of this mixture distribution:

$$\hat{\mu}_t = p_1 E\big(TN(\mu_t;\ \bar{\bar{\mu}}_t, \bar{\bar{\sigma}}_t, \alpha, \beta)\big) + p_2\alpha + p_3\beta \qquad \text{(Eq. 21)}$$

where the expectation of the truncated normal distribution is computed as:

$$E\left(TN(\mu_t;\ \bar{\bar{\mu}}_t, \bar{\bar{\sigma}}_t, \alpha, \beta)\right) = \bar{\bar{\mu}}_t - \bar{\bar{\sigma}}_t \frac{\phi(\beta)-\phi(\alpha)}{\Phi(\beta)-\Phi(\alpha)} \quad \text{(Eq. 22)}$$

where $\phi()$ represents the standard normal and $\Phi()$ its cumulative probability distribution. N.b. The variance about the generative mean, ${\sigma^2}_{\mu_t}$ (see equation 3), is thus equal to the variance of the weighted mixture distribution consisting of the truncated normal and two Dirac delta functions.

Fitting the model to participants' responses

Participants were only superficially informed about the concepts of change-points and experimental noise, without mentioning specific parameter values. Nevertheless, we assume that they obtained accurate estimates of the generative mean boundaries $\alpha = -60°$ and $\beta = 60°$ and the experimental noise $\sigma_{exp} = 10°$ by paying attention to the spatial locations of the stimuli in the training blocks and main task practice block. We also assume that they were able to form an accurate estimate of the constant hazard rate, $H = \frac{1}{6}$, during the main task practice block. We thus treat those parameters as fixed. This leaves the sensory noise ($\sigma_{sens,0°}$, $\gamma_\sigma$) and sensory gain ($\gamma_\lambda$) parameters as the only free parameters of the model.

Initial analyses of the responses in the training task (single stimulus localization) indicated that the sound localization accuracy varied considerably across participants, both in terms of biases and precision as a function of azimuth. However, the audiovisual accuracy was fairly constant across participants and azimuth. So, we decided to fix the sensory noise and bias parameters for the audiovisual stimuli to $\sigma_{sens,0°,AV} = 3°$, $\gamma_{\sigma,AV} = 0$, and $\gamma_{\lambda,AV} = 1$, but we optimized these three parameters for the auditory-only stimuli per individual participant ($\sigma_{sens,0°,A}$, $\gamma_{\sigma,A}$, $\gamma_{\lambda,A}$). The parameters were optimized by means of maximum likelihood estimation. Since we cannot know what internal estimates $\lambda_{1:t}$ an individual experienced on any given trial, we used Monte

Carlo simulations (n=1000) to integrate out the unknown sensory noise. Specifically, for every stimulus ($x_{1:t}$) in a trial we randomly sampled an internal estimate using the model's generative function:

$$\lambda_t \sim \gamma_\lambda N\left(\lambda_t; x_t, \left(\sigma_{sens,0^\circ} + \gamma_\sigma |x_t|\right)^2\right) \quad \text{(Eq. 23)}$$

Using those internal estimates $\lambda_{1:t}$ as input to the model, we then iteratively updated the beliefs about the generative mean (Eqs. 6-8, 10-17), and finally computed a best estimate of the last generative mean $\hat{\mu}_t$ (Eqs. 18-22) and last sound location $\hat{x}_t$ (Eq. 2). We then repeated this procedure one thousand times, each time sampling a new sequence of internal estimates $\lambda_{1:t}$ (Eq. 23). We thus obtained one thousand model-predicted last sound locations $\hat{x}_t$ for each trial.

We treated each of these predictions ($\hat{x}_t$) as an intended response location, but the actual response could be imprecise due to normally distributed response noise (e.g. motor noise), which we fixed at $\sigma_{resp} = 3°$. So, the likelihood of a particular response could be computed as the average (across simulations) probability density of the normal distributions that were centered on $\hat{x}_t$ with variance ${\sigma^2}_{resp}$, evaluated at response location $r$. Additionally, we assumed that participants' attention occasionally lapsed so that they had to guess the last sound location. We fixed the lapse rate to 1%. On a lapse, we assumed that participants randomly selected an intended response location from the a-priori stimulus distribution. Putting it all together, we computed the likelihood of a particular response $r$ as:

$$L(r) = \frac{0.99}{1000}\sum_{i=1}^{1000}\left(N\left(r; \hat{x}_{t,i}, {\sigma^2}_{resp}\right)\right) + \frac{0.01}{\beta-\alpha}\left(\Phi\left(\frac{\beta-r}{\sqrt{{\sigma^2}_{exp}+{\sigma^2}_{resp}}}\right) - \Phi\left(\frac{\alpha-r}{\sqrt{{\sigma^2}_{exp}+{\sigma^2}_{resp}}}\right)\right)$$

(Eq. 24)

where $i$ is an index of the Monte-Carlo simulations and $\Phi()$ represents the cumulative probability distribution of the standard normal distribution (as in Eq. 22).

Given a certain set of values for the model parameters ($\sigma_{sens,0°,A}$, $\gamma_{\sigma,A}$, $\gamma_{\lambda,A}$) we thus computed the likelihood for every response of a participant. We then used the sum of the log-likelihoods over trials as the objective function to maximize with the Bayesian adaptive direct search algorithm (BADS)[4]. We ran the algorithm four times with different starting values. Those initial parameter settings were obtained via a random search over one thousand parameter values drawn from within some preset plausible parameter interval. The four random draws that resulted in the largest log-likelihoods formed the starting positions of the optimization runs. Finally, we selected the optimized parameters that were associated with the largest (log-) likelihood across the four runs.

Prior uncertainty (PU) and surprisal (SU)

We used the model with the optimized parameters of each participant to obtain two latent variables for every stimulus. The first is the prior uncertainty (PU), which we defined as the standard deviation of the prior distribution conditional on no-changepoint, $p\left(\mu_t \middle| \lambda_{1:(t-1)}, cp_t = 0\right)$. Hence, the prior uncertainty is equal to the standard deviation of the preceding posterior distribution (Eq. 9):

$$PU_t = \bar{\bar{\sigma}}_{t-1} \quad \text{(Eq. 25)}$$

The second latent variable is the information theoretic quantity of surprisal (SU). This is defined as the negative logarithm of the probability density of the full prior distribution over internal estimates, evaluated at the latest observation $\lambda_t$:

$$SU_t = -\log\left(p\left(\lambda_t \middle| \lambda_{1:(t-1)}, cp_t = 0\right)(1-H) + p(\lambda_t | cp_t = 1)H\right) \qquad \text{(Eq. 26)}$$

where both likelihood functions are normal distributions (see equations 14 and 15), and H is the changepoint hazard rate (1/6).

Once again, we used Monte Carlo sampling to integrate out the unknown sensory noise. Specifically, we simulated 1000 sequences of internal estimates $\lambda_{1:t}$ per trial (Eq. 23) and used the model to compute the latent variables for each simulation. We then computed the average latent variable across all simulations per stimulus.

Likewise, we computed the model-predicted response locations for each trial and subject as the average intended response location $\hat{x}_t$ over 1000 simulations with individually optimized model parameters.
